# Establishing a Retron-Based Cytosine Base Editor for Targeted Hypermutation in *Escherichia coli*

**DOI:** 10.64898/2026.06.18.733067

**Authors:** Xinyu Shi, Yuyang Ni, Na Tian, Qingmin Ruan, Dingqi Liu, Jin He, Xun Wang

## Abstract

Current cytosine base editors (CBEs) are limited to unidirectional C to T conversions, restricting their applications. Retrons, bacterial genetic elements, encode a reverse transcriptase that generates multicopy single-stranded DNA (msDNA) by reverse transcribing specific non-coding RNA (ncRNA). This msDNA mimics Okazaki fragments during DNA replication, making retrons promising for gene editing. Here, we developed a retron-based cytosine base editor (RCBE) by fusing cytosine deaminase with reverse transcriptase (RT-CDA) within the retron system. RCBE first transcribes ncRNA, allowing RT-CDA to deaminate cytosine on the ncRNA. The modified ncRNA is then reverse transcribed into msDNA, where RT-CDA induces further cytosine deamination. This mutant msDNA introduces specific mutations into target gene sequences, enabling both C to T and G to A conversions. Using RCBE, we demonstrated accelerated molecular evolution of the *rpoB* gene in *Escherichia coli*. High-throughput sequencing confirmed that RCBE achieves a mutation rate of up to 0.2% in regions with high GC content. Our findings establish RCBE as a versatile tool, particularly suitable for directed evolution in GC-rich regions, with broad potential applications across various bacterial and eukaryotic hosts.

## Introduction

Natural evolution, driven by random mutations and selection over extended timescales, is inherently too slow to meet the demands of modern bioengineering for rapid acquisition of functional mutants.^1,2^ To address this limitation, directed evolution accelerates the process through iterative cycles of artificial mutagenesis and screening. While in vitro mutagenesis techniques (e.g., PCR-based mutagenesis) enable rapid generation of genetic diversity, they remain bottlenecked by laborious cloning and transformation steps.^3–5^ Thus, the development of in vivo mutagenesis systems capable of targeting specific genomic regions with precise control over mutation rates and minimal cytotoxicity represents a critical challenge in advancing directed evolution.^6–8^

Recent advances in deaminase-based technologies have provided promising solutions. Cytidine deaminases and adenosine deaminases catalyze targeted base conversions (C to T or A to G) in single-stranded DNA (ssDNA) or single-stranded RNA (ssRNA).^9–11^ For instance, CRISPR/dCas9 systems fused to deaminases exploit single-guide RNAs to localize edits at specific genomic sites, leveraging Cas9-induced R-loop formation to expose ssDNA for deamination.^12–16^ Alternative approaches, such as T7 RNA polymerase-based systems (e.g., MutaT7), fuse deaminases to processive transcription machinery to introduce mutations during gene transcription.^17–20^

Retrons, bacterial genetic elements encoding reverse transcriptases (RTs), offer a novel paradigm for targeted mutagenesis.^21^ These elements generate multicopy single-stranded DNA (msDNA) via RT-mediated reverse transcription of specific non-coding RNA (ncRNA) sequences, known as msr/msd. The resultant msDNA mimics Okazaki fragments during DNA replication, providing a template for high-efficiency genome editing in *Escherichia coli*.^22–28^ Recent studies have highlighted the potential of retrons as modular tools for directed evolution in prokaryotes, with the transcription of ncRNA driven by an error-prone T7 RNA polymerase fused with cytosine deaminase pmCDA1. This setup enables the introduction of in vivo random mutations into msDNA, which can be incorporated into the target sequence for applications in protein evolution.^29,30^

The ssDNA-generating mechanism of the retron system is closely aligned with the ssDNA/ssRNA-dependent activity of deaminases. Leveraging this alignment, we developed the Retron-mediated cytosine base editor (RCBE), which integrates Eco1 reverse transcriptase from the retron system with high-activity cytosine deaminases such as rAPOBEC1 and pmCDA1. The RCBE fusion protein retains the functionality of the reverse transcriptase and introduces C to U mutations in both ncRNA and msDNA. The mutated msDNA acts as an Okazaki fragment during replication, effectively incorporating the mutations into the target DNA while bypassing the need for Cas9 and the risk of DNA double-strand breaks (DSBs). Our results demonstrate that RCBE enables rapid iterative evolution of target sequences in *E. coli*. Given that retrons have been highlighted as modular tools for editing across kingdoms of life,^23^ we speculate that RCBE has the potential for broad adaptability across diverse biological systems.

## Results

### Design and mechanism of RCBE: Engineering CDA-RT fusion proteins

To implement the RCBE, we engineered a series of CDA-RT fusion proteins by combining each candidate CDA (pmCDA1 and rAPOBEC1) with the RT from Retron-Eco1. This design was tested with the RT attached to either the N-terminus or C-terminus of the CDA, with the two components linked by three different flexible amino acid linkers: a 100-amino-acid-long linker, a 16-amino-acid XTEN linker, and a 9-amino-acid (GGS)□ linker,^12,31,32^ as shown in Figure 1A. To enhance mutation efficiency, we also fused an uracil DNA glycosylase inhibitor (UGI) from *Bacillus subtilis* bacteriophage PBS1 to the C-terminus of each fusion protein using a short 4-amino-acid SGGS linker (Figure 1A). UGI prevents the cellular repair mechanisms from excising U, the initial product of cytosine deamination, stabilizing the intended mutation.^12^

**Figure 1.**
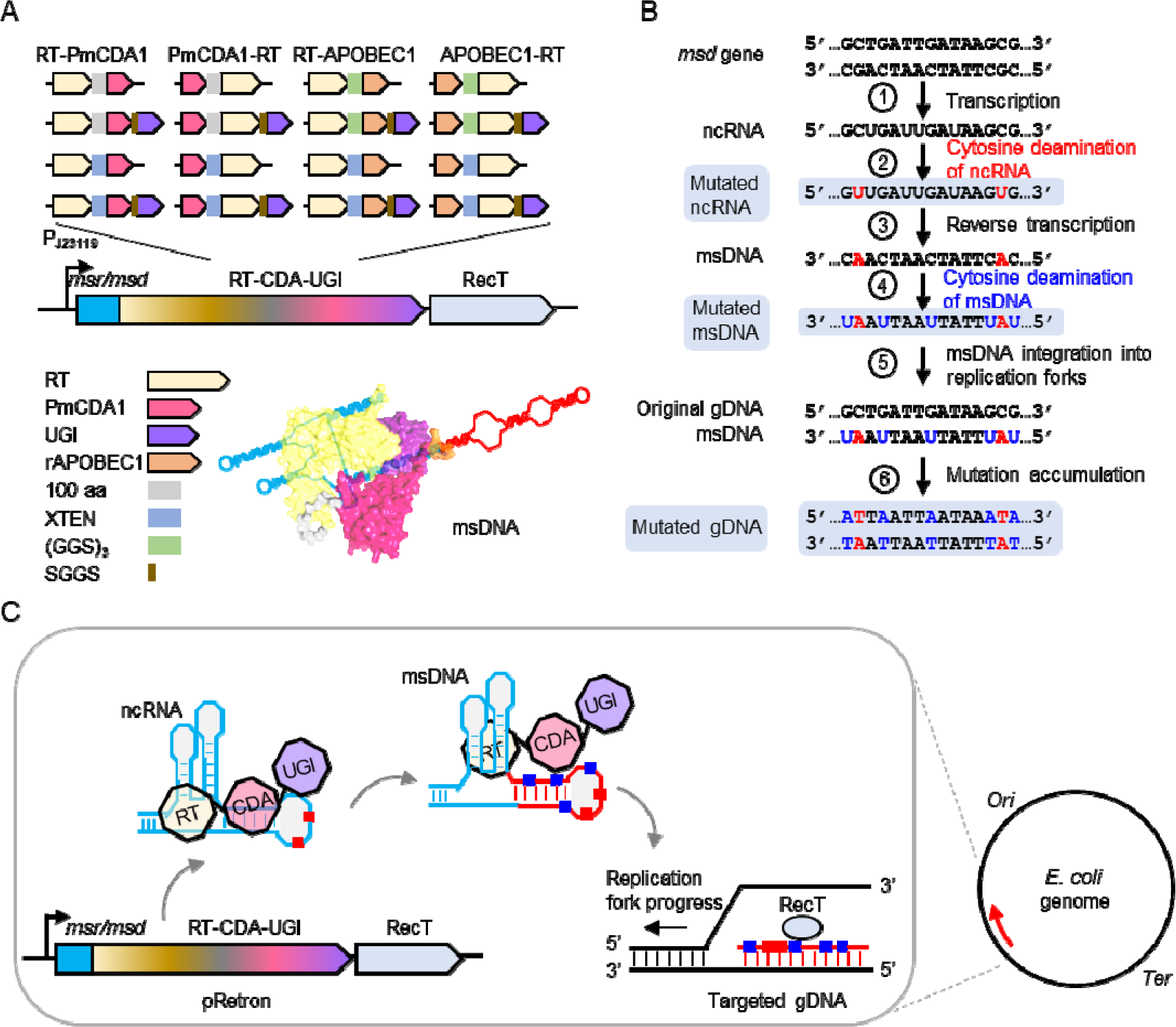
Schematic of the principle and mechanism of RCBE. (A) Schematic of the fusion protein design: Reverse transcriptase (RT) is fused to the N– or C-terminus of cytosine deaminases (PmCDA1 or rAPOBEC1) via a flexible linker. A DNA glycosylase inhibitor (UGI) is fused to the C-terminus of each fusion protein through an SGGS linker. Under the control of the strong constitutive promoter J23119, the retron operon, which includes sequences for msr/msd, the coding gene for the fusion protein, and the single-stranded DNA recombinase RecT, is expressed. On the bottom: a schematic of the RT-CDA-UGI complex with msDNA. The RT-CDA-UGI fusion protein recognizes ncRNA and reverse transcribes it to generate msDNA. (B) RCBE-mediated base editing through deamination of ncRNA and msDNA: The *msd* template is transcribed into ncRNA, where C is deaminated to U by CDA. Through reverse transcription, the mutated ncRNA is converted to msDNA, and the corresponding sequence in msDNA is mutated to A. Meanwhile, C in msDNA is deaminated to U by CDA. The mutated msDNA binds to the target DNA, replacing the Okazaki fragment during replication. After several rounds of replication, C and G on the non-template DNA are converted to T and A. (C) Schematic of RCBE-mediated targeted mutagenesis process: The *msr/msd* genes are transcribed into ncRNA, which is then recognized and bound by the RT component, and subsequently reverse transcribed into msDNA. During DNA replication, the mutated msDNA, acting as a mutation carrier, integrates mutations into the target genomic DNA under the action of RecT, thereby completing genome editing. The red box indicates deamination occurring on the ncRNA, while the blue box indicates deamination on the msDNA. These deamination events accumulate mutations in the msDNA, which ultimately triggers mutations in the target sequence.

Given that CDA can deaminate both ssRNA and ssDNA, we propose the following mechanism for RCBE-mediated mutation at the target site: Initially, the *msd* gene is transcribed to generate ncRNA (step 1 in Figure 1B). Subsequently, the fusion protein specifically recognizes and binds to the target ncRNA, exposing the ssRNA region to CDA, which deaminates C in ssRNA to U (step 2 in Figure 1B). The RT component of the fusion protein then recognizes the ncRNA and reverse transcribes the msd region, producing msDNA (step 3 in Figure 1B). CDA further deaminates C in msDNA to U (step 4 in Figure 1B). The mutated msDNA integrates into the lagging strand of the DNA replicon as an Okazaki fragment (step 5 in Figure 1B). After multiple rounds of DNA replication, the original C-G/G-C base pairs in the target gene are gradually converted to T-A/A-T base pairs, leading to the accumulation of mutations (step 6 in Figure 1B). It is worth noting that if the deamination occurs on ncRNA, the mutation at the target site is C to T. If it occurs on msDNA, the mutation is G to A.

### Verification of RCBE activity in gene editing

To ascertain the efficacy of RCBE in inducing mutations in target genes, it was imperative to verify that the RT component of our fusion protein preserved its capability to perform reverse transcription and generate msDNA for gene editing purposes. To this end, we devised a kanamycin-sensitive reporter system.

Initially, we engineered a plasmid incorporating the neomycin phosphotransferase II gene, *nptII*, which confers kanamycin resistance (Kan^R^). We subsequently introduced two premature stop codons (L23* and Y24*) within the *nptII* sequence, yielding a kanamycin-sensitive reporter plasmid designated as pKan^R^OFF (Figure 2A). Complementing this, we constructed an additional plasmid, pRetron, harboring the retron operon. This operon constitutively expresses msDNA(Kan^R^)ON, which encompasses a 75-base pair homologous sequence to the wild-type *nptII* gene. This homology facilitates the potential reversion of premature stop codons within the *nptII* gene through homologous recombination, thus restoring kanamycin resistance upon successful editing (Figure 2A).

**Figure 2.**
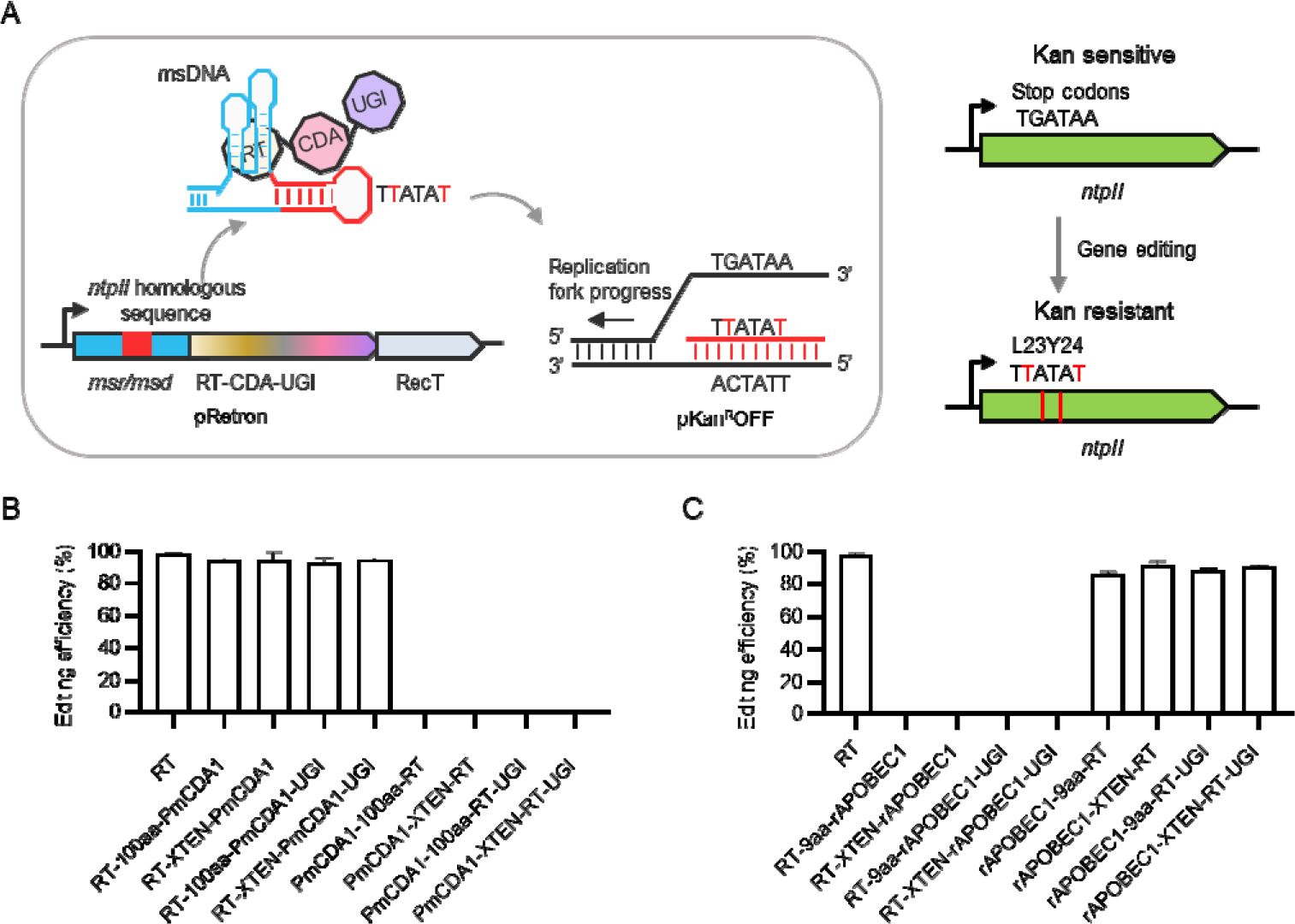
Assessment of RCBE editing efficiency. (A) The Δ*mutSrecJsbcB* strain was transformed with both pRetron and pKan^R^OFF plasmids. pRetron drives the expression of msDNA, which targets the *nptII* gene on the pKan^R^OFF plasmid. Mutation of the TGATAA sequence within the *nptII* gene to TTATAT confers kanamycin resistance to the strain. (B) Editing efficiency of strains expressing various RT, PmCDA1, and UGI fusion proteins. (C) Editing efficiency of strains expressing different RT, rAPOBEC1, and UGI fusion proteins. Data are presented as bar graphs, with each bar denoting the mean value, and error bars representing the standard deviation. The experiments were conducted with three biological replicates (n = 3).

We generated a set of 16 distinct pRetron plasmids, each encoding a unique RT-CDA fusion protein configuration as delineated in Figure 1A. These plasmids were introduced into a Δ*mutSrecJsbcB*-pKan^R^OFF strain. Post a 24-hour incubation period, we assessed the editing efficiency. Our findings revealed that successful gene editing via RCBE was attainable exclusively when PmCDA1 or the PmCDA1-UGI fusion was appended to the C-terminus of RT (Figure 2B) or when rAPOBEC1 or the rAPOBEC1-UGI fusion was attached to the N-terminus of RT (Figure 2C). Although the RCBE constructs incorporating CDA exhibited a modest decrease in editing efficiency when compared to the deaminase-lacking retron control (referred to as RT), the efficiency remained notably above 86% (Figures 2B, C). Guided by these observations, we selected RCBE constructs that maintained gene editing activity for further validation in subsequent mutation effect studies.

### Confirmation of deoxyuridine-mediated gene mutation

To substantiate the capacity of deoxyuridine present on msDNA to instigate mutations within the target gene, we chemically synthesized a 75-base pair oligonucleotide, designated as Oligo UT. In this construct, the 33rd nucleotide was specifically designed as deoxyuridine, while the 37th was thymidine (T), with the remaining sequence complementing the *nptII* gene (Figure 3A). To augment the oligonucleotide’s stability, phosphorothioate bonds were incorporated at both the 5′ and 3′ termini^33,34^. The hypothesis was that if deoxyuridine effectively mediates gene editing, it would prompt the reversion of the premature stop codon sequence TGATAA in the *nptII* gene back to the wild-type sequence TTATAT, thereby endowing the strain with kanamycin resistance (Figure 3A). As a positive control, we also synthesized Oligo TT, which features thymidine at both the 33rd and 37th positions, with the rest of the sequence being identical to that of Oligo UT.

**Figure 3.**
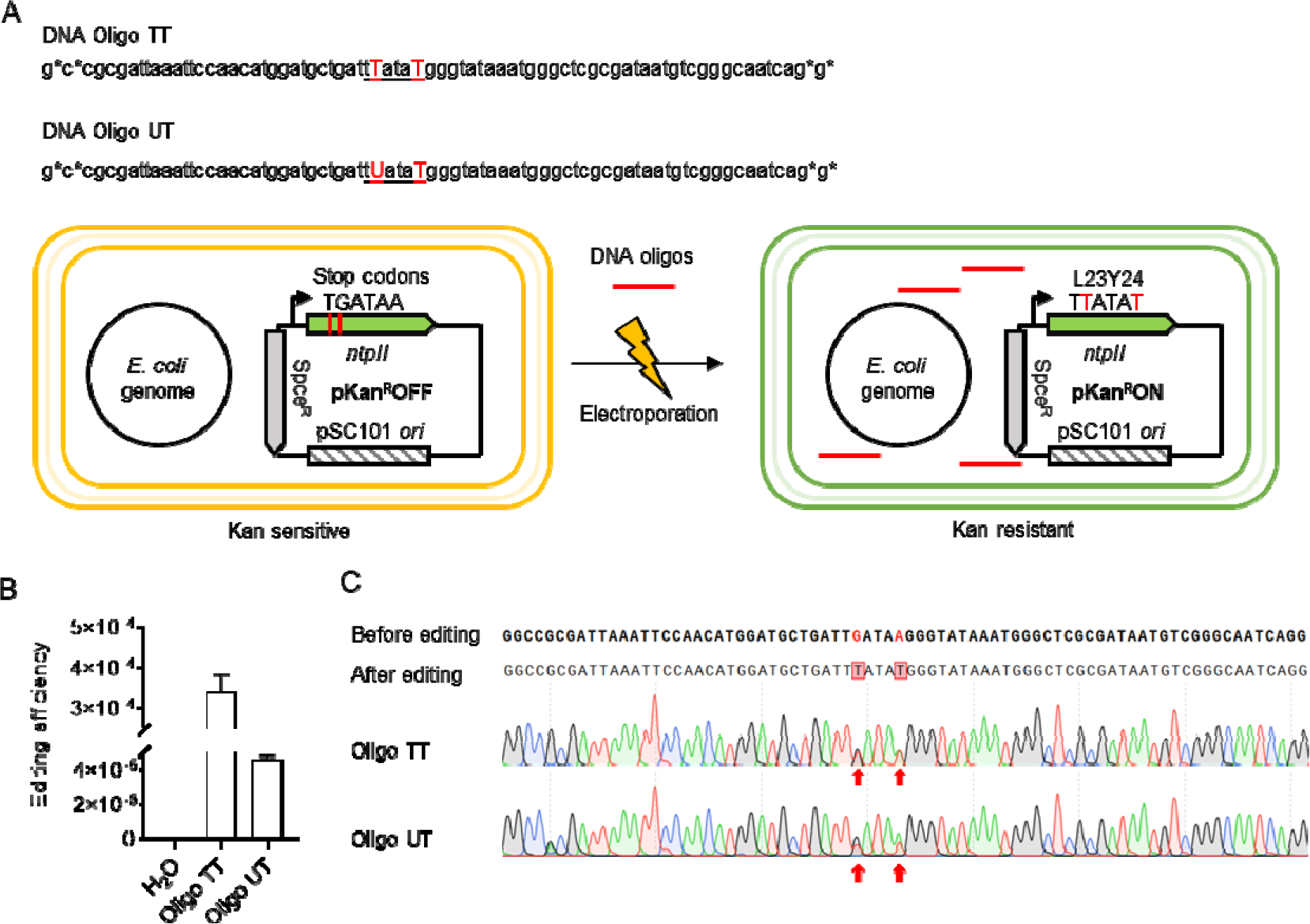
Gene editing utilizing synthesized ssDNA oligonucleotides. (A) Two ssDNA oligonucleotides with phosphorothioate-capped ends were designed. The sequences homologous to the *nptII* gene are shown in standard font, with mutations highlighted in red uppercase letters. These oligonucleotides were introduced into the Δ*mutSrecJsbcB*-pKan^R^OFF strain via electroporation. Successful editing of the *nptII* gene converts the premature stop codon TGATAA to the wild-type TTATAT, restoring kanamycin resistance. (B) Mutation efficiency assessment: Bar graph showing the proportion of kanamycin-resistant colonies resulting from the electroporation of HLJO, oligo TT, and oligo UT into the Δ*mutSrecJsbcB*-pKan^R^OFF strain. Each bar represents the mean value, and error bars indicate the standard deviation. Three biological replicates (n = 3) were performed to ensure the reliability of the results. (C) Sanger sequencing verification: Sanger sequencing results confirming the mutation at the target site. The mutated sequences are indicated by red arrows.

To evaluate the efficiency of mutation induction, Oligo TT and Oligo UT were electroporated into the Δ*mutSrecJsbcB*-pKan^R^OFF strain. As a negative control, electroporation of HDO into the Δ*mutSrecJsbcB*-pKan^R^OFF strain was performed and is referred to as the HDO control. The mutation rate was subsequently determined. Our findings revealed that the mutation efficiency for Oligo TT was 3.8 × 10DD, whereas for Oligo UT, it was 4.3 × 10DD (Figure 3B). Sanger sequencing of the selected clones confirmed the occurrence of G to T and A to T mutations at the targeted sites across all colonies, thereby validating the efficacy of deoxyuridine in mediating gene mutation (Figure 3C).

### RCBE-induced mutations in the targeted gene

To evaluate the base-editing capabilities of the RCBE system, we integrated the aminoglycoside adenylyltransferase gene A (*aadA*) into a plasmid. Specifically, we replaced the start codon (ATG) of *aadA* with nine consecutive ACG triplets, rendering the strain susceptible to streptomycin. Our hypothesis was that targeted deamination to correct any of these ACGs back to ATG would restore the expression of aminoglycoside adenylyltransferase, thereby conferring streptomycin resistance to the strain.

To assess the base-editing efficiency of RCBE on both ncRNA and msDNA, we constructed two distinct plasmids, pStrep^R^OFF1 and pStrep^R^OFF2. In pStrep^R^OFF1, the open reading frame (ORF) of the *aadA* gene was oriented clockwise, with the target sequence for msDNA being ACG, requiring the complementary sequence on msDNA to be CGT. Deamination occurring on the ncRNA was the only mechanism that could mutate the target ACG to ATG (Figure 4A). In contrast, pStrep^R^OFF2 had the *aadA* gene’s ORF oriented counterclockwise, with the targeted sequence as CGT, necessitating the complementary sequence on msDNA to be ACG. Here, deamination on msDNA was the sole mechanism capable of mutating the target ACG to ATG (Figure 4B). Thus, pStrep^R^OFF1 and pStrep^R^OFF2 plasmids respectively reflect deamination events on ncRNA and msDNA.

**Figure 4.**
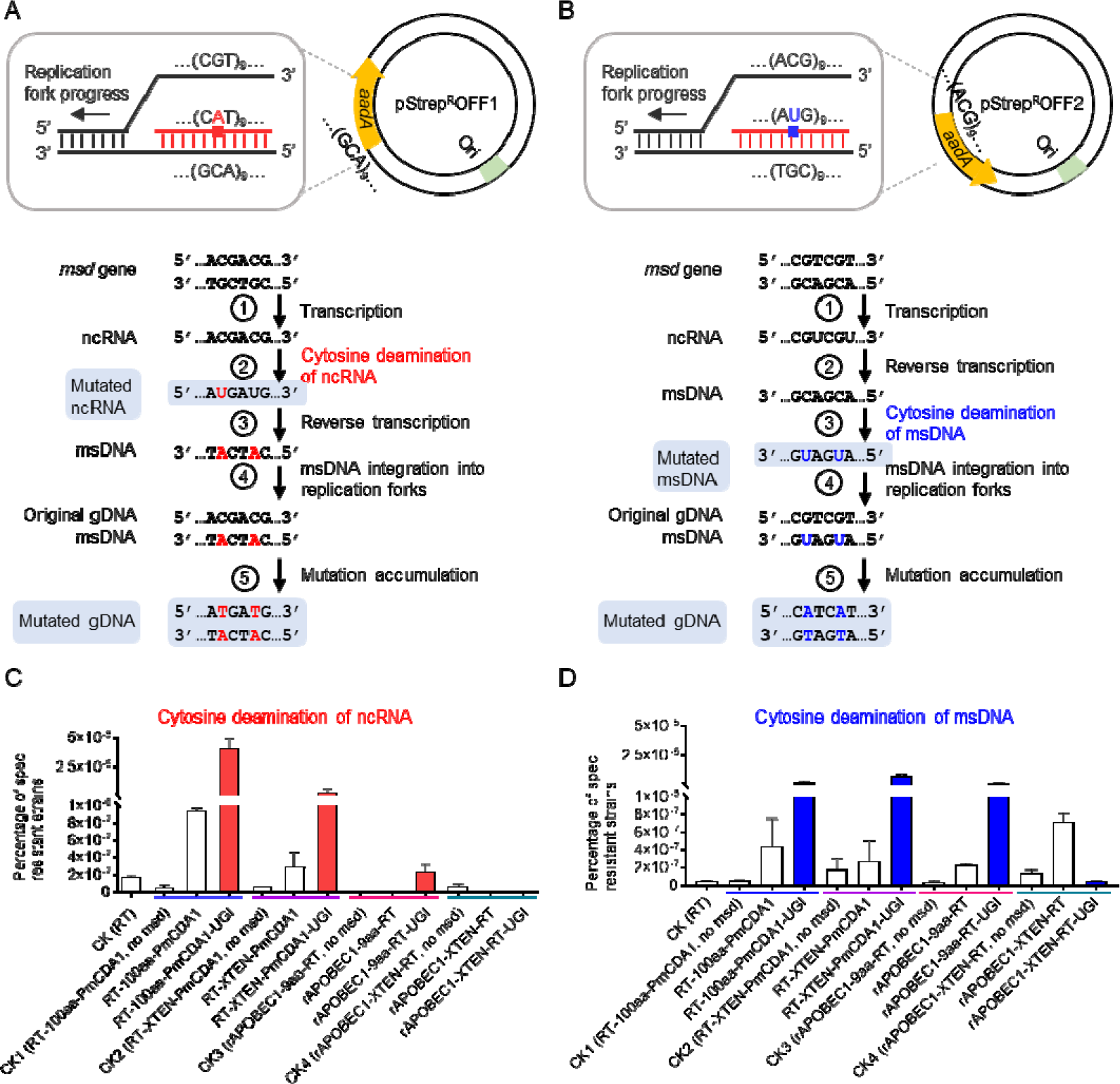
RCBE-mediated targeted mutations and their verification. (A) Schematic of RCBE-induced mutation on ncRNA: The RCBE complex, containing RT, CDA, and UGI, binds to the ncRNA. The CDA component deaminates C to U within the ncRNA sequence, leading to a mutation. The mutated ncRNA sequence is highlighted in red. (B) Schematic of RCBE-induced mutation on msDNA: The msDNA is generated by the RT component of the RCBE complex from the ncRNA. The CDA component then deaminates C to U in the msDNA sequence. The mutated msDNA sequence is highlighted in blue. (C) Bar graph illustrating the mutation efficiency of various RCBE variants on ncRNA: Each bar depicts the mean mutation rate, with error bars representing the standard deviation, based on three biological replicates (n=3). (D) Bar graph depicting the mutation efficiency of different RCBE variants on msDNA: Each bar shows the mean mutation rate, and the error bars indicate the standard deviation, based on three biological replicates (n=3).

We then constructed eight variants of RCBE that maintain editing efficiency and target the *aadA* gene: RT-100aa-PmCDA1, RT-100aa-PmCDA1-UGI, RT-XTEN-PmCDA1, RT-XTEN-PmCDA1-UGI, rAPOBEC1-9aa-RT, rAPOBEC1-9aa-RT-UGI, rAPOBEC1-XTEN-RT, and rAPOBEC1-XTEN-RT-UGI, along with four negative control variants (CK1 to CK4) containing deleted *msd* sequences. These twelve RCBE systems each carry two distinct msDNA sequences, enabling them to target the ACG regions of the *aadA* gene on pStrep^R^OFF1 and pStrep^R^OFF2, respectively. The Δ*mutSrecJsbcB*-pStrep^R^OFF strains were transformed with each of these twelve plasmids. After a 24-hour incubation, we counted the number of streptomycin-resistant colonies and calculated the mutation efficiency as the percentage of streptomycin-resistant colonies relative to the total number of colonies.

Our results indicated that among the strains characterized by deamination of C on ncRNA with the pStrep^R^OFF1 plasmid, RT-100aa-PmCDA1-UGI exhibited the highest mutation rate, reaching 4×10^-5, which is 720 times higher than the mutation rate of the control without msDNA (CK1) (Figure 4C). The mutation rate of RT-100aa-PmCDA1 was 16.8 times higher than CK1 (Figure 4C). In contrast, the mutation rates of RT-XTEN-PmCDA1 and RT-XTEN-PmCDA1-UGI were only 5.1 times and 50 times higher than CK2, respectively (Figure 4C). For RCBE variants containing rAPOBEC1, the mutation rates of rAPOBEC1-9aa-RT and rAPOBEC1-9aa-RT-UGI increased by 0.5 times and 27 times compared to CK3, respectively (Figure 4C). However, the mutation rates of rAPOBEC1-XTEN-RT and rAPOBEC1-XTEN-RT-UGI were only 0.01 times and 0.2 times higher than CK4 (Figure 4C).

Among the strains characterized by deamination of C on msDNA with the pStrep^R^OFF2 plasmid, RT-100aa-PmCDA1-UGI, RT-XTEN-PmCDA1-UGI, and rAPOBEC1-9aa-RT-UGI all demonstrated relatively high mutation rates, around 2×10^-6, which is about 40 times higher than their respective control strains CK1, CK2, and CK3 (Figure 4D). In contrast, the RCBE system containing rAPOBEC1-XTEN-RT-UGI showed no significant deamination activity (Figure 4D).

Subsequently, we selected streptomycin-resistant colonies and sequenced their *aadA* gene. Sequencing results revealed that most streptomycin resistance mutations occurred at the 1st, 2nd, and 9th ACG loci, where ACGs were corrected back to ATGs. The mutation at the 1st ACG was the most frequent (Supplementary Figure S1), likely due to its proximity to the Shine-Dalgarno sequence. Additionally, we observed a C to T mutation (mutant 2) and a G to A mutation (mutant 4) in the non-ACG region of the homologous sequence. These findings confirm that RCBE induces C to T and G to A mutations in the target region.

### Second-generation sequencing validation of RCBE-induced targeted mutations

To comprehensively evaluate the mutagenic capabilities of the RCBE system, especially in high GC-content regions, we introduced a GC-rich DNA sequence into a plasmid and subjected it to RCBE-mediated evolution. This approach allowed us to assess the system’s performance in regions where traditional selection methods, such as streptomycin resistance, may fail to detect all induced mutations. Specifically, mutations within the ACG region, such as G mutations or combined C and G mutations, do not confer resistance to spectinomycin and are thus not selected for. Therefore, we employed high-throughput sequencing (HTS) to provide an unbiased assessment of the mutation sequences and rates.

To accurately evaluate the mutation sequences and rates, we performed HTS on strains transformed with CK1, RT-100aa-PmCDA1, RT-100aa-PmCDA1-UGI, CK2, RT-XTEN-PmCDA1, and RT-XTEN-PmCDA1-UGI. The strains were cultured for 24 hours prior to sequencing the target region.

We focused on C to T and G to A mutations at the target site, which are the theoretically possible mutations induced by the RCBE system (as depicted in Figure 1B). To assess the specificity of the RCBE system, we also analyzed G-to-C mutations, which would be unexpected if mutations were random. The HTS results indicate that mutation rates for G to A and C to T were significantly higher in strains containing the RT-100aa-PmCDA1-UGI and RT-XTEN-PmCDA1-UGI variants compared to control strains (CK) (Figure 5A, B). Notably, the mutation rate for RT-100aa-PmCDA1-UGI reached up to 0.2% in the targeted region. In contrast, the occurrence of non-expected mutations, such as G-to-C mutations, was significantly lower than the expected mutations and showed no correlation with the presence of the deaminase (Figure 5C). These findings confirm that the RCBE system effectively induces targeted mutations at C and G sites, converting them to T and A, respectively.

**Figure 5.**
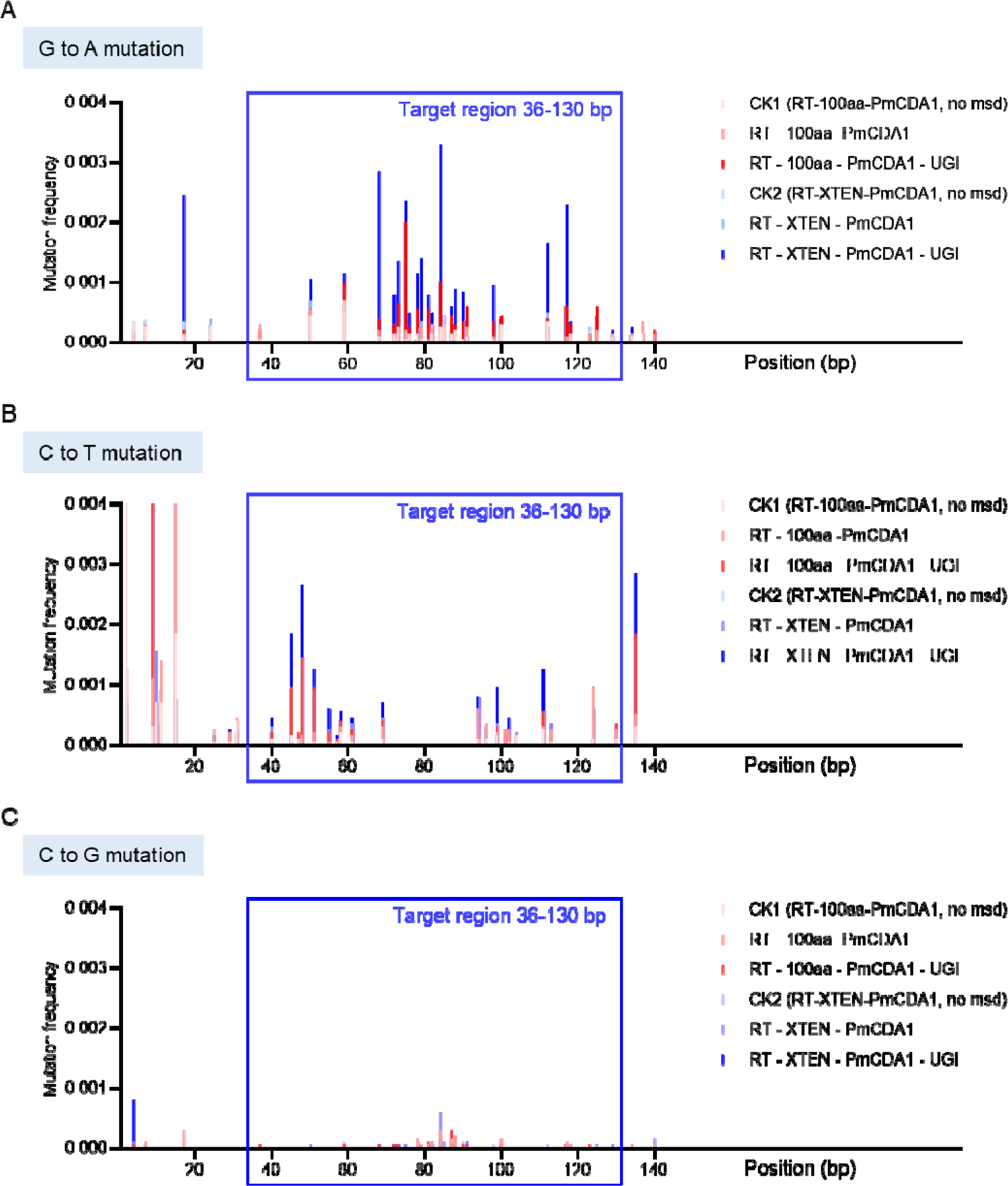
Analysis of mutation rates induced by RCBE in the targeted region. (A-C) Bar graphs depicting mutation rates for G to A, C to T, and C to G transitions, respectively, within the 94 bp homologous region targeted by msDNA, highlighted with a blue box in each panel.

### RCBE-mediated directed evolution of the *rpoB* gene in *E. coli*

Rifampicin’s bactericidal action involves binding to the β-subunit of RNA polymerase (encoded by *rpoB*), inhibiting transcription and causing cell death^35^. Mutations in *rpoB*, particularly within the rifampicin resistance-determining region (RRDR), reduce rifampicin binding and confer resistance^36,37^.

To evaluate the mutation potential of the RCBE system in practical applications, we developed an RCBE system targeting the RRDR region of the *rpoB* gene. We hypothesized that mutations induced by the RCBE system would increase the frequency of rifampicin resistance emergence in *E. coli*. We constructed two RCBE systems, F1 and F2, with homology sequence lengths of 82 bp and 84 bp, respectively, covering the coding regions for amino acids 501–528 and 529–556 in the RRDR region. As controls, we also constructed plasmids CK1 (no msd), CK2 (F1 no CDA), CK3 (F2 no CDA), and CK4 (a Δ*mutSrecJsbcB* strain), which lack the homologous sequence or the CDA component.

We cultured the strains in liquid medium for 12 and 24 hours, followed by serial dilution and spotting on solid medium containing rifampicin to quantify the number of rifampicin-resistant bacteria. The results indicated that the proportion of rifampicin-resistant strains in F1 and F2 was significantly higher than that in the control strains at both 12 and 24 hours (Figure 6A), demonstrating the effectiveness of the RCBE system for in vivo mutation and directed evolution. Notably, strains lacking the deamination system (CK2 and CK3) showed a higher proportion of resistant strains compared to CK1 or CK4, suggesting that mutation rates are elevated in msDNA-targeted regions even in the absence of deamination (Figure 6A).

**Figure 6.**
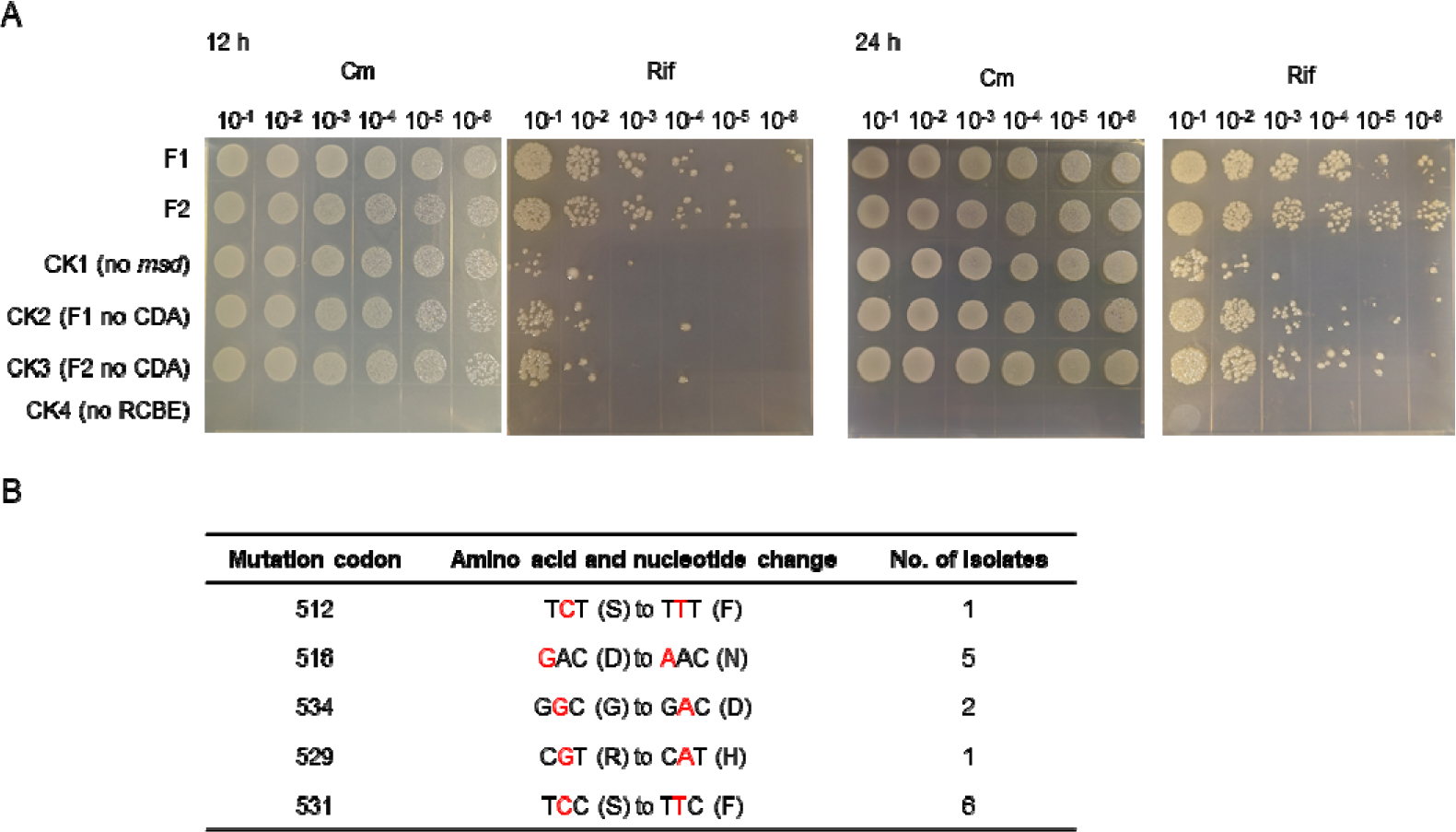
Evaluation of RCBE-mediated rifampicin resistance in *E. coli*. (A) Plate assay for rifampicin resistance: Agar plates showing bacterial growth at different dilutions after 12 and 24 hours of incubation. The plates contain rifampicin (Rif) and chloramphenicol (Cm) to select for transformants. (B) Mutation analysis of rifampicin resistance-conferring sites: Table summarizing the mutations identified in the RRDR of the *rpoB* gene that confer rifampicin resistance. The mutations are listed by their codon number, the amino acid and nucleotide change, and the number of isolates exhibiting each mutation.

We then randomly selected 20 rifampicin-resistant strains and sequenced the RRDR region. Among them, 15 strains displayed the expected G-to-A or C-to-T mutations, all involving single-point mutations. The mutation at serine 531 was the most prevalent, observed in six strains, followed by a mutation at aspartic acid 516 (Figure 6B). Mutations were also identified at amino acid positions 512, 534, and 529. The remaining five strains exhibited random mutations unrelated to the deamination system, which we speculate may be due to the inherently high mutation rate of the RT enzyme^38^. These findings suggest that the RCBE system is highly effective in inducing targeted gene mutations.

## Discussion

In this study, we present the development of a novel retron-based cytosine base editor (RCBE), which utilizes cytidine deaminase (CDA) as a mutation-inducing tool. The system harnesses the unique transcription and reverse transcription mechanisms of retrons to generate msDNA, which serves as a “mutation carrier” to induce C-to-T and G-to-A mutations at target sequences in vivo.

One of the standout features of RCBE is its remarkable flexibility in target sequence selection. Unlike other gene editing tools, RCBE can introduce mutations in virtually any region of interest within the genome. This broad target sequence adaptability enhances its versatility, making it suitable for a wide array of gene editing applications. Furthermore, RCBE is capable of inducing both C-to-T and G-to-A mutations, providing a broader range of mutation types, which is particularly valuable for directed evolution in GC-rich regions and the creation of genetic variants.

RCBE offers several distinct advantages over traditional methods such as lambda recombineering or ssDNA oligonucleotide pools. First, it requires minimal exogenous DNA, as retrons generate msDNA in vivo, streamlining the process. Second, RCBE provides a more efficient in vivo mutagenesis method by bypassing the need for complex plasmid library construction, which is often required for oligonucleotide pool methods. This makes RCBE especially advantageous for rapid, direct mutations in bacterial genomes. Additionally, its ability to facilitate iterative mutations at a single genetic locus is a powerful tool for directed evolution, enabling the generation of diverse variants with greater efficiency than conventional techniques.

Looking ahead, several improvements could further enhance the RCBE system. Co-expressing CDA with adenosine deaminase could enable simultaneous mutations across all four DNA bases (A, T, C, and G), thereby increasing mutation diversity. Additionally, optimizing the retron system-such as modifying the Retron-Eco1 system or exploring other retrons that can accommodate longer DNA fragments could extend the mutational window and improve editing efficiency. Slowing the activity of RT may also increase the interaction time between RT, ncRNA, and msDNA, boosting mutation efficiency without compromising msDNA production.

In summary, RCBE represents a promising new approach for gene editing and directed evolution, offering key advantages in target sequence flexibility, mutation diversity, and iterative mutagenesis. With further optimization, RCBE has the potential to become a crucial tool in genetic research and biotechnology.

## Materials and Methods

### Bacterial strains and growth conditions

*E. coli* DH5α was used for plasmid construction. The Δ*mutSrecJsbcB* strain was employed to assess both mutation rates and editing efficiency^28^. *E. coli* strains carrying the corresponding plasmids (as indicated in the figures) were cultured at 37°C in Lysogeny Broth (LB) medium containing 10 g of tryptone, 5 g of yeast extract, and 10 g of NaCl per liter of water, and supplemented with kanamycin (50 μg/mL), chloramphenicol (34 μg/mL), streptomycin (50 μg/mL), or rifampicin (100 μg/mL), as required.

### Plasmid construction

Primers were designed using SnapGene software, with lengths typically ranging from 15 to 30 bp and G+C content between 40% and 60%. The primers used for plasmid construction were synthesized by Tianyi Huiyuan (Wuhan, Hubei, China). All plasmids generated in this study were assembled using the Hieff Clone Plus Multi One-Step Cloning Kit (Yeasen, Shanghai, China). Briefly, insert and vector fragments were amplified by PCR and then assembled into the vector through homologous recombination. The resulting plasmids were validated by sequencing (Quintarabio, Wuhan, Hubei, China). These plasmids were subsequently introduced into bacterial cells using the calcium chloride (CaCl□) method to produce the corresponding derivative strains. Details of the coding sequences for genetic elements used in this study (e.g., cytidine deaminase, resistance genes, protein linkers) are provided in Supplementary Table S1.

### Editing efficiency testing

For plasmid-based editing, plasmids containing various RT-CDAs were introduced alongside the pKan^R^OFF plasmid into Δ*mutSrecJsbcB* competent cells. Briefly, 100 µL of the competent cells were incubated with 100 pmol plasmids on ice for 20 minutes, followed by heat-shocking at 42°C for 90 seconds and rapid cooling on ice for 2 minutes. The cells were then supplemented with 900 µL of LB medium and incubated at 37°C for 1 hour. Following this incubation, 200 µL of the culture was added to 5 mL of LB medium containing chloramphenicol and streptomycin, and the mixture was incubated for 24 hours. The culture was then diluted to 10□□ and plated onto LB agar plates containing either chloramphenicol and streptomycin or chloramphenicol and kanamycin. Editing efficiency was calculated as the ratio of colony counts on the kanamycin plates to those on the streptomycin plates.

For DNA oligo-mediated editing, 100 pmol of oligonucleotides TT and UT were electroporated into 100 µL of the Δ*mutSrecJsbcB*-pKan^R^OFF cells. The cells were then supplemented with 900 µL of LB medium and incubated for 2.5 hours. After incubation, 1 µL, 10 µL, and 100 µL aliquots of the culture were plated onto LB agar plates containing chloramphenicol and kanamycin. Additionally, dilutions of 10□□, 10□□, and 10□□ were plated onto LB agar plates containing chloramphenicol and streptomycin. Editing efficiency was determined by calculating the ratio of colony counts on kanamycin plates to those on streptomycin plates.

### RCBE system mutation efficiency testing

To evaluate mutation efficiency in the RCBE system, Δ*mutSrecJsbcB*-pStrep^R^OFF competent cells were transformed with plasmids carrying various RT-CDA. After transformation, cells were incubated in LB medium with chloramphenicol and kanamycin for 1 hour at 37°C. Subsequently, 200 µL of the culture was transferred to 5 mL of LB medium with selective antibiotics, followed by a 24-hour incubation.

Aliquots of 1 µL, 10 µL, and 100 µL from the culture were then plated onto LB agar plates with chloramphenicol and streptomycin, while serial dilutions (10□□, 10□□, and 10□□) were plated onto plates containing chloramphenicol and kanamycin. Mutation efficiency was calculated as the ratio of streptomycin-resistant to kanamycin-resistant colonies.

### High-throughput DNA sequencing (HTS)

*E. coli* strains harboring the specified RCBE systems were cultured for 24 hours, after which target sequences were amplified using high-fidelity polymerases to ensure accurate replication. The amplified DNA products were then sent for sequencing (Tsingke, Wuhan, Hubei, China). Upon receiving the raw sequencing data, sequence pairs were assembled with FLASH v1.2.11 software, followed by alignment of the assembled sequences to the reference template using BLASTN for precise mutation analysis. Python scripts were subsequently employed to calculate single-nucleotide mutation frequencies at each base position, providing detailed insights into mutation distribution and efficiency.

## Acknowledgments

This work was supported by the National Key Research and Development Program of China (2022YFF1000700), the Hubei Provincial Major Special Project for Industrial Development of Agricultural Microorganisms (NYWSWZX2025-2027-10) and the Hubei Provincial Natural Science Foundation of China (JCZRLH202600436).

## Conflict of interest

The authors declare that they have no conflict of interest.

## Notes

### Competing Interest Statement

The authors have declared no competing interest.

## References

1 Packer, M. S. & Liu, D. R. Methods for the directed evolution of proteins. Nature Reviews Genetics 16, 379–394 (2015). 10.1038/nrg3927

2 Castle, S. D., Stock, M. & Gorochowski, T. E. Engineering is evolution: a perspective on design processes to engineer biology. Nature Communications 15 (2024). https://doi.org/ARTN 3640 10.1038/s41467-024-48000-1

3 Yuan, L., Kurek, I., English, J. & Keenan, R. Laboratory-Directed Protein Evolution. Microbiology and Molecular Biology Reviews 69, 373–392 (2005). 10.1128/mmbr.69.3.373-392.2005

4 Fujii, R., Kitaoka, M. & Hayashi, K. Error-prone rolling circle amplification: the simplest random mutagenesis protocol. Nat Protoc 1, 2493–2497 (2006). 10.1038/nprot.2006.403

5 Zhao, H., Giver, L., Shao, Z., Affholter, J. A. & Arnold, F. H. Molecular evolution by staggered extension process (StEP) in vitro recombination. Nature Biotechnology 16, 258–261 (1998). 10.1038/nbt0398-258

6 Romero, P. A. & Arnold, F. H. Exploring protein fitness landscapes by directed evolution. Nature Reviews Molecular Cell Biology 10, 866–876 (2009). 10.1038/nrm2805

7 Wang, Y. J. et al. Directed Evolution: Methodologies and Applications. Chem Rev 121, 12384–12444 (2021). 10.1021/acs.chemrev.1c00260

8 Cao, M. F., Tran & Zhao, H. M. Unlocking nature’s biosynthetic potential by directed genome evolution. Curr Opin Biotech 66, 95–104 (2020). 10.1016/j.copbio.2020.06.012

9 Hodges, P. E., Navaratnam, N., Greeve, J. C. & Scott, J. Site-specific creation of uridine from cytidine in apolipoprotein B mRNA editing. Nucleic acids research 19, 1197–1201 (1991). 10.1093/nar/19.6.1197

10 Conticello, S. G. The AID/APOBEC family of nucleic acid mutators. Genome Biol 9, 229 (2008). 10.1186/gb-2008-9-6-229

11 Pecori, R., Di Giorgio, S., Paulo Lorenzo, J. & Nina Papavasiliou, F. Functions and consequences of AID/APOBEC-mediated DNA and RNA deamination. Nature reviews. Genetics 23, 505–518 (2022). 10.1038/s41576-022-00459-8

12 Komor, A. C., Kim, Y. B., Packer, M. S., Zuris, J. A. & Liu, D. R. Programmable editing of a target base in genomic DNA without double-stranded DNA cleavage. Nature 533, 420–424 (2016). 10.1038/nature17946

13 Wang, Y. et al. In-situ generation of large numbers of genetic combinations for metabolic reprogramming via CRISPR-guided base editing. Nature Communications 12, 678 (2021). 10.1038/s41467-021-21003-y

14 Ma, Y. et al. Targeted AID-mediated mutagenesis (TAM) enables efficient genomic diversification in mammalian cells. Nat Methods 13, 1029–1035 (2016). 10.1038/nmeth.4027

15 Halperin, S. O. et al. CRISPR-guided DNA polymerases enable diversification of all nucleotides in a tunable window. Nature 560, 248–252 (2018). 10.1038/s41586-018-0384-8

16 Gaudelli, N. M. et al. Programmable base editing of A•T to G•C in genomic DNA without DNA cleavage. Nature 551, 464–471 (2017). 10.1038/nature24644

17 Moore, C. L., Papa, L. J., 3rd & Shoulders, M. D. A Processive Protein Chimera Introduces Mutations across Defined DNA Regions In Vivo. J Am Chem Soc 140, 11560–11564 (2018). 10.1021/jacs.8b04001

18 Mengiste, A. A. et al. Expanded MutaT7 toolkit efficiently and simultaneously accesses all possible transition mutations in bacteria. Nucleic Acids Research 51, e31–e31 (2023). 10.1093/nar/gkad003

19 Park, H. & Kim, S. Gene-specific mutagenesis enables rapid continuous evolution of enzymesin vivo. Nucleic Acids Research 49, e32–e32 (2021). 10.1093/nar/gkaa1231

20 Álvarez, B., Mencía, M., de Lorenzo, V. & Fernández, L. Á. In vivo diversification of target genomic sites using processive base deaminase fusions blocked by dCas9. Nature Communications 11 (2020). 10.1038/s41467-020-20230-z

21 Simon, A. J., Ellington, A. D. & Finkelstein, I. J. Retrons and their applications in genome engineering. Nucleic acids research 47, 11007–11019 (2019). 10.1093/nar/gkz865

22 Farzadfard, F. & Lu, T. K. Genomically encoded analog memory with precise in vivo DNA writing in living cell populations. 346, 1256272 (2014). doi:10.1126/science.1256272

23 Lopez, S. C., Crawford, K. D., Lear, S. K., Bhattarai-Kline, S. & Shipman, S. L. Precise genome editing across kingdoms of life using retron-derived DNA. Nat Chem Biol 18, 199–206 (2022). 10.1038/s41589-021-00927-y

24 Ellis, H. M., Yu, D., DiTizio, T. & Court, D. L. High efficiency mutagenesis, repair, and engineering of chromosomal DN A using single-stranded oligonucleotides. Proceedings of the National Academy of Sciences 98, 6742–6746 10.1073/pnas.121164898

25 Kong, X. et al. Precise genome editing without exogenous donor DNA via retron editing system in human cells. Protein & cell 12, 899–902 (2021). 10.1007/s13238-021-00862-7

26 Schubert, M. G. et al. High-throughput functional variant screens via in vivo production of single-stranded DNA. Proceedings of the National Academy of Sciences of the United States of America 118 (2021). 10.1073/pnas.2018181118

27 Sharon, E. et al. Functional Genetic Variants Revealed by Massively Parallel Precise Genome Editing. Cell 175, 544–557.e516 (2018). 10.1016/j.cell.2018.08.057

28 Ni, Y. et al. Reducing competition between msd and genomic DNA improves retron editing efficiency. EMBO reports 25, 5316–5330 (2024). 10.1038/s44319-024-00311-6

29 Liu, W. et al. Retron-mediated multiplex genome editing and continuous evolution in Escherichia coli. Nucleic acids research 51, 8293–8307 (2023). 10.1093/nar/gkad607

30 Simon, A. J., Morrow, B. R. & Ellington, A. D. Retroelement-Based Genome Editing and Evolution. Acs Synth Biol 7, 2600–2611 (2018). 10.1021/acssynbio.8b00273

31 Schellenberger, V. et al. A recombinant polypeptide extends the in vivo half-life of peptides and proteins in a tunable manner. Nat Biotechnol 27, 1186–1190 (2009). 10.1038/nbt.1588

32 Wang, Y. et al. MACBETH: Multiplex automated Corynebacterium glutamicum base editing method. Metab Eng 47, 200–210 (2018). 10.1016/j.ymben.2018.02.016

33 Wang, H. H. et al. Programming cells by multiplex genome engineering and accelerated evolution. Nature 460, 894–898 (2009). 10.1038/nature08187

34 Isaacs, F. J. et al. Precise manipulation of chromosomes in vivo enables genome-wide codon replacement. Science 333, 348–353 (2011). 10.1126/science.1205822

35 Wehrli, W., Knusel, F., Schmid, K. & Staehelin, M. Interaction of rifamycin with bacterial RNA polymerase. Proc Natl Acad Sci U S A 61, 667–673 (1968). 10.1073/pnas.61.2.667

36 Goldstein, B. P. Resistance to rifampicin: a review. J Antibiot (Tokyo) 67, 625–630 (2014). 10.1038/ja.2014.107

37 Jin, D. J. & Gross, C. A. Mapping and sequencing of mutations in the Escherichia coli rpoB gene that lead to rifampicin resistance. J Mol Biol 202, 45–58 (1988). 10.1016/0022-2836(88)90517-7

38 Jamburuthugoda, V. K. & Eickbush, T. H. The reverse transcriptase encoded by the non-LTR retrotransposon R2 is as error-prone as that encoded by HIV-1. J Mol Biol 407, 661–672 (2011). 10.1016/j.jmb.2011.02.015

