## Supplemental Figure 1 and Supplemental Table 1 for "Establishing a Retron-Based Cytosine Base Editor for Targeted Hypermutation in *Escherichia coli*": Supplementary Files.docx

**Table of contents**

Figure S1

Sanger sequencing results of the translation start region of the *aadA* gene	P. 2

Table S1

Coding sequences for genetic elements used in this study	P. 3

**Supplementary Figure S1**

Sanger sequencing results of the translation start region of the *aadA* gene, with the mutated sequences marked by red arrows.
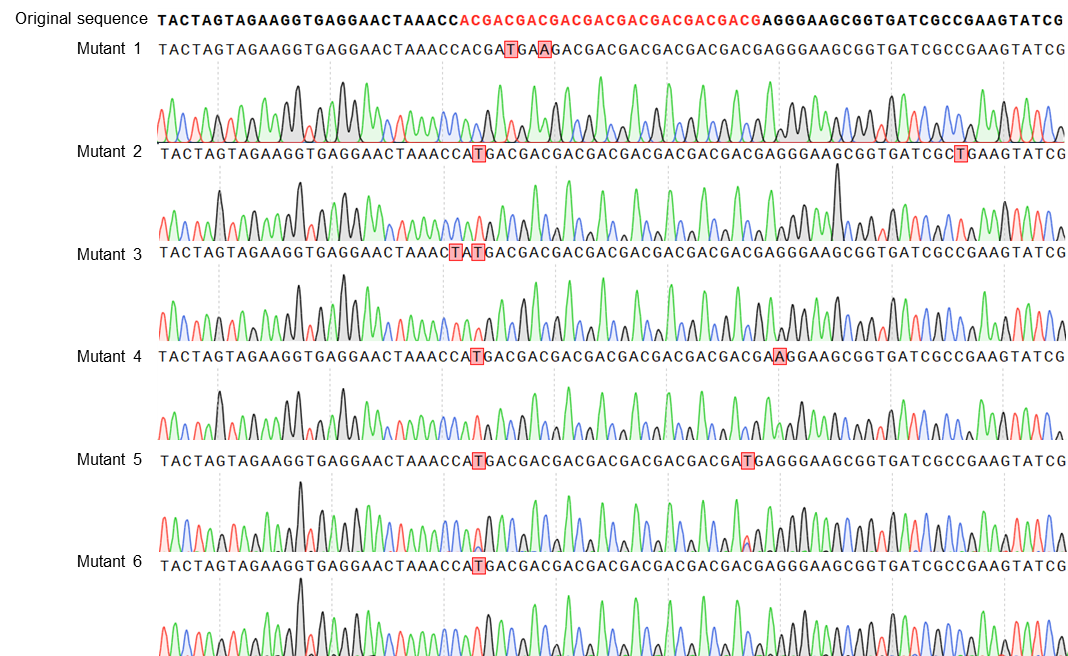

**Table S1 Coding sequences for genetic elements used in this study**

| **Component name** | **Function** | **Sequence 5’ to 3’** |
| --- | --- | --- |
| PmCDA1 | Cytidine Deaminase,  introduces C-to-T (or G-to-A) mutations | ATGACAGATGCGGAATATGTGAGAATTCATGAAAAACTGGATATCTATACGTTCAAAAAGCAGTTTTTCAACAACAAAAAGAGCGTGAGCCATAGATGCTATGTGCTGTTCGAACTGAAAAGAAGAGGAGAAAGAAGAGCGTGCTTTTGGGGCTATGCGGTGAATAAACCGCAGAGCGGAACGGAAAGAGGAATTCATGCGGAAATTTTTAGCATTAGAAAAGTCGAAGAATACCTGAGAGATAATCCGGGACAATTTACAATTAATTGGTATAGCTCTTGGAGCCCGTGCGCGGATTGCGCAGAAAAAATTCTGGAATGGTATAATCAGGAACTGAGAGGCAATGGCCATACACTGAAAATTTGGGCGTGCAAACTGTATTATGAAAAAAATGCGAGAAATCAGATTGGCCTGTGGAATCTGAGAGATAATGGCGTCGGCCTGAATGTCATGGTCAGCGAACATTATCAGTGCTGCAGAAAAATTTTTATTCAGAGCAGCCATAATCAGCTGAATGAAAATAGATGGCTGGAAAAAACACTGAAAAGAGCGGAAAAAAGAAGAAGCGAACTGAGCATTATGATTCAGGTCAAAATTCTGCATACAACAAAAAGCCCGGCGGTC |
| APOBEC-1 |  | ATGTCATCAGAAACCGGACCAGTTGCAGTTGATCCAACCCTGAGACGGCGGATAGAACCACACGAATTTGAAGTTTTTTTCGACCCTCGTGAACTTAGAAAAGAAACCTGTCTGCTGTATGAAATTAATTGGGGAGGTCGTCATAGTATTTGGCGCCATACCAGCCAGAACACCAATAAACATGTTGAAGTTAATTTCATCGAGAAGTTTACCACCGAACGCTATTTTTGTCCCAACACCCGCTGCAGCATTACATGGTTCCTGAGCTGGAGTCCTTGCGGAGAATGCAGCCGGGCCATTACCGAATTTCTGTCTCGCTATCCGCATGTTACCCTGTTTATTTATATTGCACGTCTGTATCATCACGCCGACCCGCGTAATCGGCAGGGGCTGCGTGATCTGATTTCATCTGGTGTTACCATTCAGATTATGACCGAACAAGAAAGTGGCTATTGTTGGCGTAACTTTGTGAATTATTCTCCTAGTAATGAGGCTCATTGGCCAAGATATCCTCATCTTTGGGTTCGTCTGTATGTTCTGGAACTGTATTGCATTATCCTGGGTCTGCCGCCGTGTTTAAATATTCTGCGTCGCAAGCAACCGCAGTTAACCTTTTTTACCATTGCCCTGCAATCTTGTCATTATCAACGGCTGCCGCCTCATATTCTGTGGGCCACCGGTTTAAAA |
| UGI | Inhibits uracil-DNA glycosylase to prevent uracil excision | ACCAATTTAAGTGATATTATTGAAAAGGAGACCGGCAAACAGCTGGTTATCCAGGAAAGCATTTTAATGCTGCCTGAAGAAGTTGAAGAAGTTATTGGAAATAAGCCGGAAAGCGATATCCTGGTTCATACCGCCTATGATGAAAGCACCGACGAAAACGTTATGCTGTTAACCTCCGACGCACCGGAATATAAACCGTGGGCACTGGTGATTCAGGACAGCAATGGTGAAAATAAAATTAAAATGCTG |
| Linker-100aa | Links protein domains in fusion proteins | CCGAAAAAGAAAAGAAAAGTCGGCGGCGGCGGAAGCGGAGGAGGAGGATCAGCAGAATATGTCAGAGCGCTGTTTGATTTTAATGGCAATGACGAAGAAGACCTGCCGTTTAAAAAGGGCGACATTCTGAGAATTAGAGATAAACCGGAAGAACAGTGGTGGAATGCGGAAGATAGCGAAGGAAAAAGAGGCATGATTCCGGTACCGTATGTCGAAAAATACTCAGGCGATTATAAAGATCATGATGGAGATTATAAAGACCATGATATTGATTACAAAGATGACGATGATAAAAGCAGA |
| Linker-SGGS |  | AGCGGTGGTAGC |
| Linker-XTEN |  | AGCGGATCAGAAACACCGGGAACCTCAGAATCAGCAACCCCGGAAAGC |
| Linker-GGS×3 |  | GGCGGCAGCGGTGGTAGCGGCGGCAGC |
| *aadA* | aminoglycoside adenylyltransferase gene A (*aadA*), the start codon (ATG) of *aadA* was replaced with nine consecutive ACGs, rendering the strain susceptible to streptomycin. | TAATAATTGACAGCTAGCTCAGTCCTAGGTATAATACTAGTAGAAGGTGAGGAACTAAACCACGACGACGACGACGACGACGACGACGAGGAGGGAAGCGGTGATCGCCGAAGTATCGACTCAACTATCAGAGGTAGTTGGCGTCATCGAGCGCCATCTCGAACCGACGTTGCTGGCCGTACATTTGTACGGCTCCGCAGTGGATGGCGGCCTGAAGCCACACAGTGATATTGATTTGCTGGTTACGGTGACCGTAAGGCTTGATGAAACAACGCGGCGAGCTTTGATCAACGACCTTTTGGAAACTTCGGCTTCCCCTGGAGAGAGCGAGATTCTCCGCGCTGTAGAAGTCACCATTGTTGTGCACGACGACATCATTCCGTGGCGTTATCCAGCTAAGCGCGAACTGCAATTTGGAGAATGGCAGCGCAATGACATTCTTGCAGGTATCTTCGAGCCAGCCACGATCGACATTGATCTGGCTATCTTGCTGACAAAAGCAAGAGAACATAGCGTTGCCTTGGTAGGTCCAGCGGCGGAGGAACTCTTTGATCCGGTTCCTGAACAGGATCTATTTGAGGCGCTAAATGAAACCTTAACGCTATGGAACTCGCCGCCCGACTGGGCTGGCGATGAGCGAAATGTAGTGCTTACGTTGTCCCGCATTTGGTACAGCGCAGTAACCGGCAAAATCGCGCCGAAGGATGTCGCTGCCGACTGGGCAATGGAGCGCCTGCCGGCCCAGTATCAGCCCGTCATACTTGAAGCTAGACAGGCTTATCTTGGACAAGAAGAAGATCGCTTGGCCTCGCGCGCAGATCAGTTGGAAGAATTTGTCCACTACGTGAAAGGCGAGATCACCAAGGTAGTCGGCAAATAA |
| Retron *msd* Kan^R^ | Editing the *nptII* gene in pKan^R^OFF | GCCGCGATTAAATTCCAACATGGATGCTGATTTATATGGGTATAAATGGGCTCGCGATAATGTCGGGCAATCAGG |
| Retron *msd* Strep^R^ | Editing the *aadA* gene in pStrep^R^OFF | ATAATACTAGTAGAAGGTGAGGAACTAAACCACGACGACGACGACGACGACGACGACGAGGGAAGCGGTGATCGCCGAAGTATCGACTCAAC |
